# Neolithic and Medieval virus genomes reveal complex evolution of Hepatitis B

**DOI:** 10.1101/315531

**Authors:** Ben Krause-Kyora, Julian Susat, Felix M. Key, Denise Kühnert, Esther Bosse, Alexander Immel, Christoph Rinne, Sabin-Christin Kornell, Diego Yepes, Sören Franzenburg, Henrike O. Heyne, Thomas Meier, Sandra Lösch, Harald Meller, Susanne Friederich, Nicole Nicklisch, Kurt Werner Alt, Stefan Schreiber, Andreas Tholey, Alexander Herbig, Almut Nebel, Johannes Krause

## Abstract

The hepatitis B virus (HBV) is one of the most widespread human pathogens known today, yet its origin and evolutionary history are still unclear and controversial. Here, we report the analysis of three ancient HBV genomes recovered from human skeletons found at three different archaeological sites in Germany. We reconstructed two Neolithic and one medieval HBV genomes by *de novo* assembly from shotgun DNA sequencing data. Additionally, we observed HBV-specific peptides using paleo-proteomics. Our results show that HBV circulates in the European population for at least 7000 years. The Neolithic HBV genomes show a high genomic similarity to each other. In a phylogenetic network, they do not group with any human-associated HBV genome and are most closely related to those infecting African non-human primates. These ancient virus forms appear to represent distinct lineages that have no close relatives today and went possibly extinct. Our results reveal the great potential of ancient DNA from human skeletons in order to study the long-time evolution of blood borne viruses.

## Introduction

The hepatitis B virus (HBV) is one of the most widespread human pathogens, with one third of the world population being infected, and an annual death toll of about 1 million globally (WHO, 2017). Infection of liver cells with HBV leads to acute hepatitis B, which is self-limiting in about 90-95% of cases. In about 5-10% of infected individuals virus clearance fails and patients develop chronic infection of hepatitis B, which puts them at lifelong elevated risk for liver cirrhosis and liver cancer (hepatocellular carcinoma). HBV is usually transmitted by contact with infectious blood, in highly endemic countries often during birth (WHO, 2017).

HBV has a circular, partially double-stranded DNA genome of about 3.2kbp that encodes four overlapping open reading frames (P, pre-S/S, pre-C/C, and X). Based on the genomic sequence diversity, HBVs are currently classified into 8 genotypes (A-H) and numerous subgenotypes that show distinct geographic distributions (Castelhano et al., 2017). All genotypes are hypothesised to be primarily the result of recombination events (Littlejohn et al., 2016; Simmonds and Midgley, 2005). To a lesser extent, HBV evolution is also driven by the accumulation of point mutations (Schaefer 2007, Araujo 2015).

Despite being widespread and well-studied, the origin and evolutionary history of HBV is still unclear and controversial (Littlejohn et al., 2016, Souza et al., 2014). HBVs in non-human primates (NHP), for instance in chimpanzees and gorillas, are phylogenetically closely related to, and yet distinct from, human HBV isolates, supporting the notion of an Africa origin of the virus (Souza et al., 2014). Molecular-clock based analyses dating the origin of HBV have resulted in conflicting estimates with some as recent as about 400 years ago (Zhou and Holmes, 2007, Souza et al., 2014). These observations have raised doubts about the suitability of molecular dating approaches for reconstructing the evolution of HBV (Bouckaer et al., 2103, Souza et al., 2014). Moreover, ancient DNA (aDNA) research on HBV-infected mummies from the 16^th^ century AD revealed a very close relationship between the ancient and modern HBV genomes (Kahila Bar-Gal et al., 2012, Patterson Ross et al., 2018), indicating a surprising lack of temporal genetic changes in the virus during the last 500 years (Patterson Ross et al., 2018). Therefore, diachronic aDNA HBV studies, in which both the changes in the viral genome over time as well as the provenance and age of the archaeological samples, are needed to better understand the origin and evolutionary history of the virus.

Here, we report the analysis of three complete HBV genomes recovered from human skeletal remains from the prehistoric Neolithic and Medieval Periods in Central Europe. Our results show that HBV already circulated in the European population more than 7000 years ago. Although the ancient forms show a relationship to modern isolates they appear to represent distinct lineages that have no close modern relatives and are possibly extinct today.

## Results and Discussion

We detected evidence for presence of ancient HBV in three human tooth samples as part of a metagenomic screening for viral pathogens that was performed on shotgun sequencing data from 53 skeletons using the metagenomic alignment software MALT (Vagene et al., 2018). The remains of the individuals were excavated from the Neolithic sites of Karsdorf (Linearbandkeramik [LBK], 5056–4959 cal BC) and Sorsum (Tiefstichkeramik group of the Funnel Beaker culture, 3335-3107 cal BC), the medieval cemetery of Petersberg/Kleiner Madron (1020-1116 cal AD), all located in Germany (Fig. A, figure supplementary S1-S3). After the three aDNA extracts had appeared HBV-positive in the initial virus screening, they were subjected to deep-sequencing without any prior enrichment resulting in 367 to 419 million reads per sample (table 1). Analysis of the human DNA recovered from Karsdorf (3-fold coverage), Sorsum (1.2-fold coverage) and Petersberg (2.9-fold coverage) showed genetic affinities of the individuals to LBK, Funnel Beaker from Sweden and medieval human populations, respectively (figure supplementary S9-S11), which is in agreement with the archeological evidence. The results of the human population genetic investigation as well as the typical aDNA deamination patterns in the recovered human and HBV sequences (supplementary, figure supplementary S4-S5) support the ancient origin of the obtained dataset.

**Fig A.**
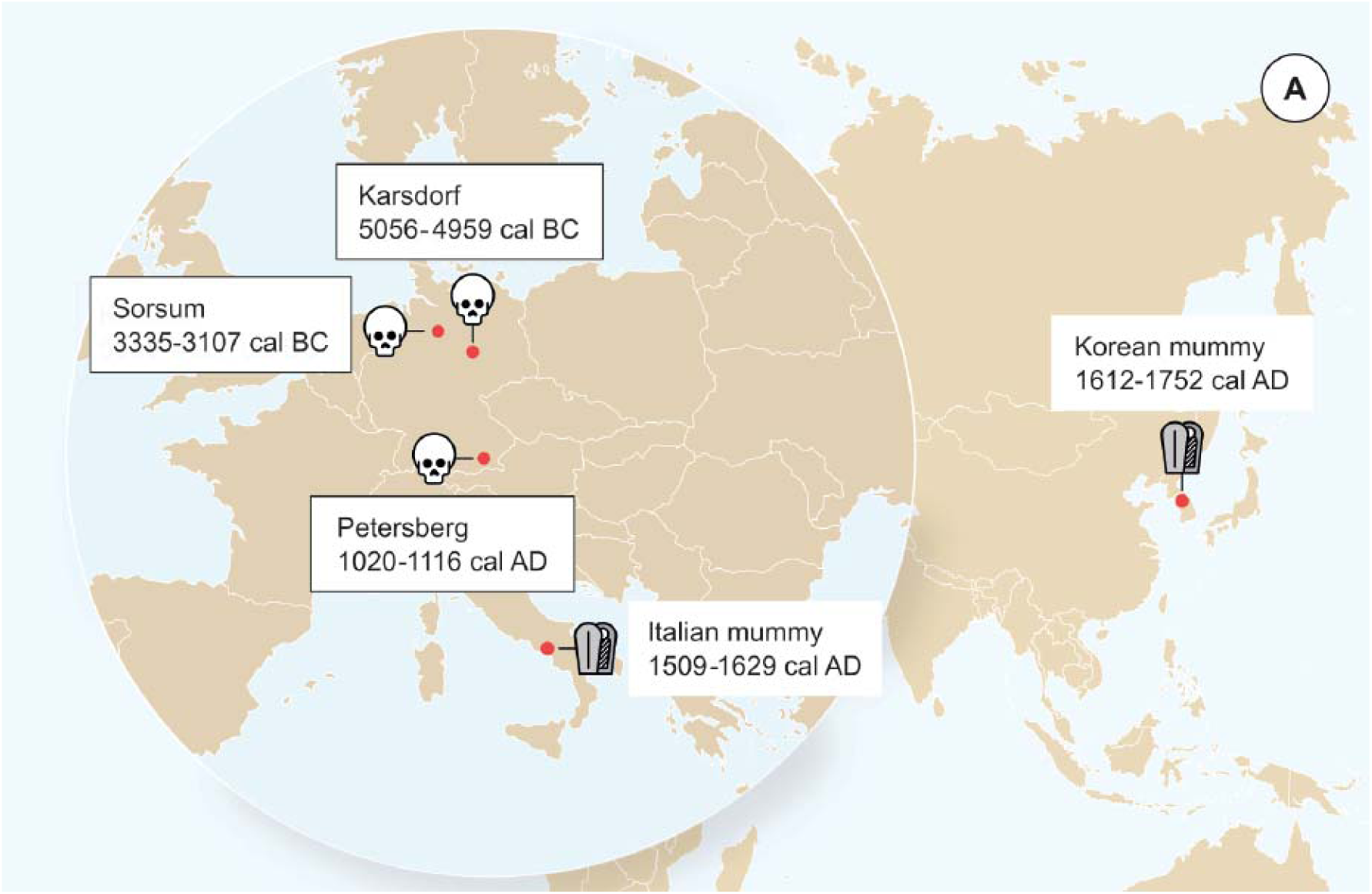
Origin of samples. Geographic location of the samples from which ancient HBV genomes where isolated. Radiocarbon dates of the specimens is given in 2 sigma range. Icon indicate the sample material (tooth or mummy). HBV genomes obtained in this study indicated by black frame.

**Table 1.** Results of the genome reconstruction.

|  | # Merged reads | Length of HBV consensus sequence | mean HBV coverage | Gaps in the consensus sequence at nt position | # mapped reads HBV | # mapped reads human | mean human coverage | human genomes/ HBV genomes |
| --- | --- | --- | --- | --- | --- | --- | --- | --- |
| **Karsdorf** | 386,780,892 | 3183 | 104X | 2157-2175;<br>3107-3128;<br>3133-3183 | 10,718 | 122,568,310 | 2.96X | 1 : 35.1 |
| **Sorsum** | 367,574,767 | 3182 | 47X | - | 3,249 | 9,856,001 | 1.17X | 1 : 40.2 |
| **Petersberg** | 419,413,082 | 3161 | 46X | 880-1000;<br>1232-1329;<br>1331-1415;<br>1420-1581;<br>1585-1598 | 2,125 | 105,476,677 | 2.88X | 1 : 16 |

For successful HBV genome reconstruction, we mapped all metagenomic sequences to 16 HBV reference genomes (8 human genotypes (A-H) and 8 NHPs from Africa and Asia) that are representative of the current HBV strain diversity (supplementary, table supplementary S6). The mapped reads were used for a *de novo* assembly, resulting in contigs from which one ancient HBV consensus sequence per sample was constructed. The consensus genomes are 3161 (46-fold coverage), 3182 (47-fold coverage), and 3183 (105-fold coverage) nucleotides in length, which falls in the length range of modern HBV genomes and suggests that we successfully reconstructed the entire ancient HBV genomes (table 1, figure supplementary S6-S8). Further, when we conducted liquid chromatography-mass spectrometry (LC-MS) based bottom-up proteomics on tooth material from the three individuals, we identified in the Karsdorf and Petersberg samples a peptide that is part of the very stable HBV core protein, supporting the presence and active replication of HBV in the individuals’ blood (supplementary, figure supplementary S16).

Phylogenetic network analysis was carried out with a dataset comprised of 495 modern HBV strains representing the full genetic diversity. Strikingly, the Neolithic HBV genomes did not group with any human strain in the phylogeny. Instead, they branched off in two clades and were most closely related to the African non-human primates (NHP) genomes (Fig. B, 93% similarity). Although the two Neolithic strains were recovered from humans who had lived about two thousand years apart, they showed a higher genomic similarity to each other than to any other human or NHP genotype. Still, their genomes differed by 6% and may therefore be considered representatives of two separate clades. The genome from the 1000-year-old Petersberg individual clustered with modern D4 genotypes.

**Fig B.**
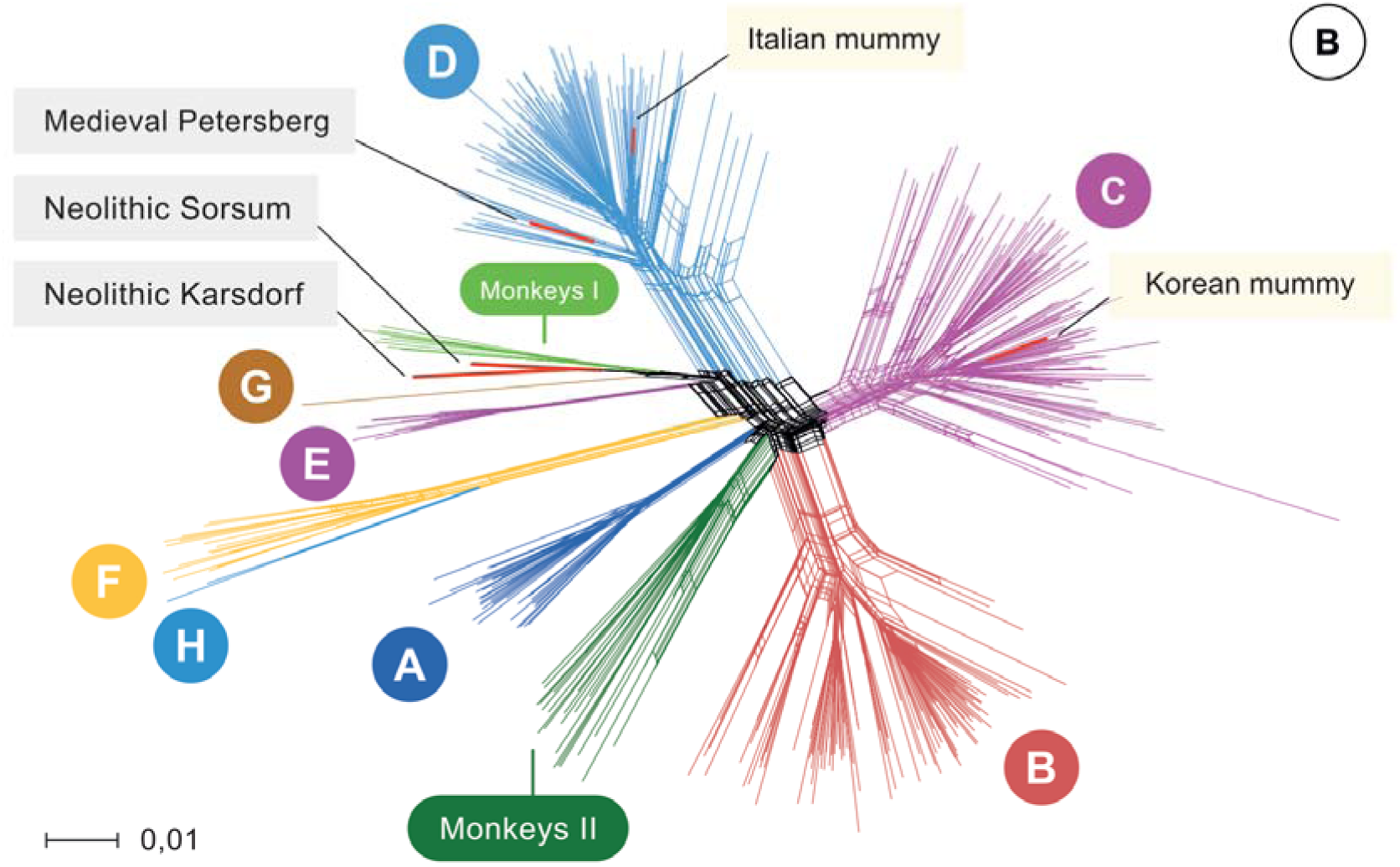
Network. Network of 495 modern, two published ancient genomes (light yellow box), and three ancient hepatitis B virus (HBV) obtained in this study (grey box). Colors indicating the 8 human HBV Genotypes (A-H), two monkey genotypes (Monkeys I, African apes and Monkeys II, Asian monkeys) and ancient genomes (red).

Owing to continuous recombination over time, different gene segments or modules of the ancestral genomes can show up in various subsequent virus generations. Such precursors have been postulated (Simmonds and Midgley, 2005) and their existence is supported by the results of our recombination analysis (figure supplementary S12-S15, source data 1). Some fragments of the Karsdorf sequences appeared to be very similar to modern human (G, E) and African NHP genotypes, and the Sorsum genome partially showed a high similarity to the human genotypes G, E and B. (figure supplementary S12-S13, S15, source data 1). Given the close relationship between the two Neolithic virus genomes, it is also conceivable that the older HBV from Karsdorf could have been a distant source for the younger Sorsum virus (figure supplementary S12-S13, S15, source data 1). The closer relationship between the Neolithic and the NHP strains compared to other human strains is noteworthy and may have involved reciprocal cross-species transmission at one or possibly several times in the past (Simmonds and Midgley, 2005, Souza et al., 2014, Rasche et al. 2016).

Taken together, our results demonstrate that HBV already existed in Europeans 7000 years ago and that its genomic structure closely resembled that of modern hepatitis B viruses. Both Neolithic viruses fall between the present-day modern human and the known NHP diversity. Therefore, it can be hypothesized that although the two Neolithic HBV strains are no longer observed today and thus may reflect two distinct clades that went extinct, they could still be closely related to the remote ancestors of the present-day genotypes, which is supported by signs of ancient recombination events. More ancient precursors, intermediates and modern strains of both humans and NHPs need to be sequenced to disentangle the complex evolution of HBV. As this evolution is characterized by recombination and point mutations and may further be complicated by human-ape host barrier crossing (Simmonds and Midgley, 2005, Souza et al., 2014, Rasche et al. 2016), genetic dating is not expected to yield meaningful results.

Based on our analysis, HBV DNA can reliably be detected in tooth samples that are up to 7000 years old. Ancient HBV has so far only been identified in soft tissue from two 16^th^-century mummies (Kahila Bar-Gal et al., 2012, Patterson Ross et al., 2018). The aDNA analysis of HBV from prehistoric skeletons, which facilitates evolutionary studies on a deeper temporal scale, has not been described up to now. One explanation for the difficulty of a molecular HBV diagnosis in bones is that the virus infection does not leave lesions on skeletal remains that would allow researchers to select affected individuals *a priori*, as it is the case for instance for leprosy (Schuenemann et al., 2013). The diagnosis of an HBV infection in skeletal populations is purely a chance finding and is thus more probable in a large-scale screening.

Overall, HBV biomolecules seem to be well preserved in teeth: We could reconstruct three HBV genomes by *de novo* assembly from shotgun data and even observed HBV-specific peptides. The ratio of HBV genomes to the human genome in our samples was rather high and similar in all three samples (Karsdorf 35:1, Sorsum 40.2:1 and Petersberg 16:1). As there is no evidence that HBV DNA is more resistant to postmortem degradation than human DNA, the high rate of HBV compared to human DNA may reflect the disease state in the infected individuals at the time of death. High copy numbers of viral DNA in the blood of infected individuals are associated with acute HBV infection, or reactivation of chronic HBV. Thus, it seems likely that the death of the ancient individuals is related to the HBV infection, but might not be the direct cause of death as fulminant liver failure is rather rare in modern day patients. The HBV infection might have instead contributed to other forms of lethal liver failure such as cirrhosis or liver cancer.

In view of the unexpected complexity of our findings, we envisage future diachronic HBV studies that go beyond the temporal and geographic scope of our current work.

## Materials and Methods

### Human remains

The LBK settlement of **Karsdorf**, Saxony-Anhalt, Germany, is located in the valley of the river Unstrut. Between 1996-2010 systematic excavations were conducted at Karsdorf that led to the discovery of settlements and graves from the Neolithic to the Iron Age (Behnke, 2007, 2011, 2012). The LBK is represented by 24 longhouses in north-west to south-east orientation that were associated with settlement burials (Veit, 1996). The investigated individual 537 is a male with an age at death of around 25-30 years (figure supplementary S1), dated to 5056–4959 cal BC (KIA 40357 – 6116 ± 32 BP) (Brandt et al., 2014, Nicklisch, 2017).

The gallery grave of **Sorsum**, Lower-Saxony, Germany, is typologically dated to the Tiefstichkeramik (group of the Funnelbeaker culture). Sorsum is exceptional as it was built into the bedrock. During the excavations (1956-1960) of the grave chamber around 105 individuals were recovered (Claus, 1983, Czarnetzki, 1966). Individual XLVII 11 analyzed in this study is a male (figure supplementary S2) and dates to 3335-3107 cal BC (MAMS 33641 − 4501 ± 19 BP).

The medieval cemetery on the **Petersberg/Kleiner Madron**, Bavaria, Germany, lies on a hill top at 850 meters asl and 400 meters above the floor of the Inn Valley. On the eastern part of the cemetery, which is under discussion here, members of a priory were buried that was most likely established in the late 10^th^ century. Written sources document its existence from 1132 onwards (Meier, 1998). During systematic excavations (1997-2004) in the southeastern part of the churchyard 99 graves with a higher, but hardly determinable number of individuals were uncovered. The examined individual in grave 820 is a male with an age at death of around 65-70 years (Lösch, 2009 - figure supplementary S3) dating to 1020-1116 cal AD (MAMS 33642 − 982 ± 17 BP).

### DNA extraction and sequencing

The DNA extractions and pre-PCR steps were carried out in clean room facilities dedicated to aDNA research. Teeth were used for the analyses. The samples from Petersberg and Sorsum were processed in the Ancient DNA Laboratory at Kiel University and the sample from Karsdorf in the Ancient DNA Laboratory of the Max Planck Institute for the Science of Human History (MPI SHH) in Jena. All procedures followed the guidelines on contamination control in aDNA studies (Warinner et al., 2017, Key et al., 2017). The teeth were cleaned in pure bleach solution to remove potential contaminations prior to powdering. Fifty milligrams of powder were used for extraction following a silica-based protocol (Dabney et al., 2013). Negative controls were included in all steps.

From each sample, double-stranded DNA sequencing libraries (UDGhalf) were prepared according to an established protocol for multiplex high-throughput sequencing (Meyer and Kircher, 2010). Sample-specific indices were added to both library adapters via amplification with two index primers. Extraction and library blanks were treated in the same manner. For the initial screening, the library of the individual from Karsdorf was sequenced on 1/50 of a lane on the HiSeq 3000 (2x75 bp) at the MPI SHH in Jena and the libraries from Petersberg and Sorsum were sequenced on the Illumina HiSeq 4000 (2x75 bp) platform at the Institute of Clinical Molecular Biology, Kiel University, using the HiSeq v4 chemistry and the manufacturer’s protocol for multiplex sequencing. Deep-sequencing for each of the three samples was carried out on 2 lanes on the Illumina HiSeq 4000 platform at the Institute of Clinical Molecular Biology, Kiel University.

### Metagenomics data processing, screening, and analyses

The datasets for the three ancient samples comprised paired-end reads. The adapter sequences were removed and overlapping paired-end reads were merged with ClipAndMerge which is a module of the EAGER pipeline (Peltzer et al., 2016). The metagenomic viral screening was carried out using MALT (Vagene et al., 2018) and the NCBI viral RefSeq database. All three samples showed HBV-specific reads. In order to obtain all HBV related sequencing reads we mapped against a multi-fasta reference containing one representative of each genotype (A-H) and eight ape strains using BWA (Li and Durbin, 2010) (table supplementary S6). Mapped reads were extracted from the BAM file, converted to FASTQ and a *de novo* assembly using SPAdes (Bankevich et al., 2012) was carried out. Resulting contigs for each K-value where checked and the k-value that spawned the longest contigs was selected as criteria for further analysis. The contigs were re-mapped with BWA against the multi-fasta reference. The resulting alignment was visually inspected in IGV v 2.3.92 (Thorvaldsdóttir et al., 2013) to archive information about contig order and direction. Based on that information, a consensus sequence was constructed from the contigs.

We assembled a comprehensive set of reference genomes using 5497 non-recombinant genomes available at hpvdb (https://hbvdb.ibcp.fr/HBVdb/HBVdbDataset?seqtype=0) and a previously defined set of 74 ape-infecting HBV genomes. In order to reduce the actual number of genomes used for subsequent inferences but retain the full range of known HBV diversity, we clustered all sequences using UClust v 1.1.579 (Edgar et al., 2010). We extracted the centroid sequences based on a sequence identity of at least 97%, which resulted in 495 representative genomes. Those genomes together with all available ancient genomes were aligned using Geneious version 10.1.2 (Kearse et al., 2012) with a 65% similarity cost matrix, a gap open penalty of 12 and a gap extension penalty of 3. The multiple sequence alignment was stripped of any sites (columns) that had gaps in more than 95% of sequences. The complete alignment including all modern and ancient genomes is available as multi-fasta in source data 2. The alignment was used to construct a network with the software SplitsTree v4 (Huson and Bryant, 2006), creating a NeighborNet (Bryant and Moulton, 2004) with uncorrected P distances.

### Recombination analysis

We performed recombination analysis using all modern full reference genomes (n=495) and five ancient genomes used for the network analysis (see above). The methods RDP, GENECOV, Chimera, MaxChi, BootScan, SiScan, 3Seq within RDP v4 (Martin et al., 2015) with a window size of 100 nt and the parameter set to circular genome with and without outgroup reference (results are provided in source data 1) and SimPlot v 3.5.1 (Lole et al. 1999, figure supplement S12-S15) were applied to the data set.

### Human population genetic analyses

Mapping of the adapter-clipped and merged FASTQ files to the human reference genome hg19 was done using BWA (Li and Durbin, 2010) using a reduced mapping stringency of “-n 0.01” and the mapping quality parameter “q 30”. The mapped sequencing data was transformed into the *Eigenstrat* format (Price et al., 2006) and merged with a dataset of 1.233.013 SNPs (Haak et al., 2015, Mathieson et al., 2015). Using the software Smartpca (Patterson et al., 2006) the three samples and previously published ancient populations were projected onto a base map of genetic variation calculated from 32 West Eurasian populations (figure supplement S9-S11).

### Sex determination

Sex determination was assessed based on the ratio of sequences aligning to the X and Y chromosomes compared to the autosomes (Skoglund et al., 2013).

### LC-MS based bottom-up proteomics

Proteins were extracted from powdered tooth samples (50 mg) using a modified filter-aided sample preparation (FASP) protocol as previously described (Cappellini et al., 2013, Warinner et al., 2014). Samples were digested using trypsin and analyzed by LC-MS/MS. Protein identification was performed using the SequestHT (Thermo Scientific) search engine in a combined database comprising the full Swiss protein database (468,716 entries), a hepatitis B data base (7 entries) and a common contaminant list. Further details regarding the LC-MS/MS analysis and database search parameters are given in the supplementary information and figure supplementary S16.

## Acknowledgements

We are grateful to the following people and institutions for providing samples, support, and advice: Bodo Krause-Kyora, Hildegard Nelson (Referat A1 Archäologische Dokumentation, Niedersächsisches Landesamt für Denkmalpflege), Ulrike Weller (Sammlungsverwaltung Archäologie Landesmuseum Hannover) and Britta Steer for technical assistance with proteomics sample preparation.

This work was supported by the Collaborative Research Centre 1266 *Scales of Transformation*, the Excellence Cluster 306 *Inflammation at Interfaces*, the Medical Faculty of Kiel University, the Max Planck Society and the European Research Council (ERC) starting grant APGREID (to J.K.). Excavations and analysis of the archaeological site of Karsdorf were supported by the German Research Foundation (DFG) Grant of Kurt W. Alt (Al 287-7-1) and Harald Meller (Me 3245/1-1).

## Accession numbers

Raw sequence read files have been deposited at the European Nucleotide Archive under accession no. PRJEB24921

